# Let there be multifunctionality: Uncovering the criticality zoo of the AC-DC genetic circuit

**DOI:** 10.1101/2025.08.22.671785

**Authors:** Smitha Maretvadakethope, Içvara Aor, Matthias M. Fischer, Àngela Pantebre Pedrosa, Yolanda Schaerli, Ruben Perez-Carrasco

**Affiliations:** Department of Life Sciences, Imperial College London, United Kingdom; Department of Bioengineering, Imperial College London, United Kingdom; Imperial Centre for Engineering Biology, Imperial College London, United Kingdom; Department of Fundamental Microbiology, University of Lausanne, Switzerland; Institute for Theoretical Biology, Charité Universitätsmedizin Berlin and Humboldt-Universität zu Berlin, Germany; Institute of Pathology, Charité Universitätsmedizin Berlin, Germany

## Abstract

Gene regulatory networks (GRNs) govern processes such as cell fate, patterning, and adaptation. While multistability and oscillations are both common GRN dynamics in cell biology, they are typically studied and engineered in isolation. Here, we challenge this separation using the AC–DC circuit, a minimal three-gene network that merges the classical toggle switch and repressilator. Using a thermodynamic formalism and Bayesian inference, we show that even a single-inducer version of the circuit can display diverse multifunctional dynamics, including the coexistence of oscillations and multistability. In addition we explore robustness, classify emergent behaviours, and analyse critical slowing down and regime transitions. Remarkably, the AC–DC circuit can produce more than 30 topologically distinct bifurcation diagrams, challenging the classical view that network topology rigidly constrains dynamical outcomes. This flexibility enables synthetic capabilities that couple hysteresis with oscillations, critical slowing down, and reversibility. By uncovering the hidden potential of minimal genetic circuits and outlining design principles for their implementation, this work opens new directions for harnessing emergent complexity using the basic building blocks of life.

## I. INTRODUCTION

Gene regulatory networks (GRNs) direct biological features by controlling the spatial and temporal organisation of gene expression [1]. Through the dynamic interpretation of external cues and transcription factors within the network, they govern complex regulation of biological processes such as pattern formation in multicellular organisms, adaptation to environmental changes, and the coordination of both metabolism and growth. The ability of GRNs to orchestrate such complex processes relies on the nonlinear responses of the underlying transcription and translation machinery, even with very small GRNs composed of only a few genes. Examples of such dynamic behaviours include genetic oscillations [2–5], chaotic dynamics [6], and multistable gene switches [7].

Although small GRNs have been extensively studied, the full breadth of dynamic behaviours they can exhibit remains significantly underestimated. Traditionally, network topology has been viewed as the primary determinant of function; however, this picture is much more nuanced [8]. It has become increasingly clear that circuits sharing the same topology can display strikingly different dynamical functions by simply tuning biophysical parameters, a phenomenon known as multifunctionality [9–11]. Fully uncovering this dynamic repertoire is not only crucial for understanding the capabilities of small GRNs but also represents a necessary step toward achieving a mechanistic understanding of larger, genome-scale networks, which are now routinely explored through omics approaches [12–14].

Beyond their dynamical richness, multifunctional circuits offer major advantages for cellular efficiency, enabling complex genetic programmes to be implemented with minimal resource burden [15, 16]. This minimal design principle is particularly attractive in synthetic biology, where multifunctional GRNs can be engineered to achieve specific behaviours or to enable novel biotechnological applications while minimising costs to the host [17–19]. Moreover, multifunctionality offers an evolutionary advantage, allowing cells to encode a diverse range of adaptable and evolvable behaviours using a small set of components [20, 21]. Together, these aspects underscore the fundamental and applied significance of multifunctional circuits in both natural and synthetic contexts.

One striking example of a small multifunctional circuit is the classic two-gene cross-repressive motif [22, 23]. Traditionally known for encoding a genetic toggle switch, recent studies have shown that by adjusting transcription factor binding rates, this motif can also generate irreversible memory states, act as a transient signal detector, or serve as a timer that uses noise to increase robustness [24, 25]. Another particularly intriguing example – and the focus of this manuscript – is the AC-DC circuit, a three-gene GRN (see Fig. 1a) first observed in vertebrate neural tube development [26] and later in the Drosophila gap gene network [27]. Combining elements of a bistable switch with a repressilator (cyclic sequential inhibition of three genes), the AC-DC circuit can theoretically produce rich dynamical phenomena, such as multistability between oscillations and steady states that can be used to encode oscillation synchronisation or excitability.[5, 26].

**FIG. 1.**
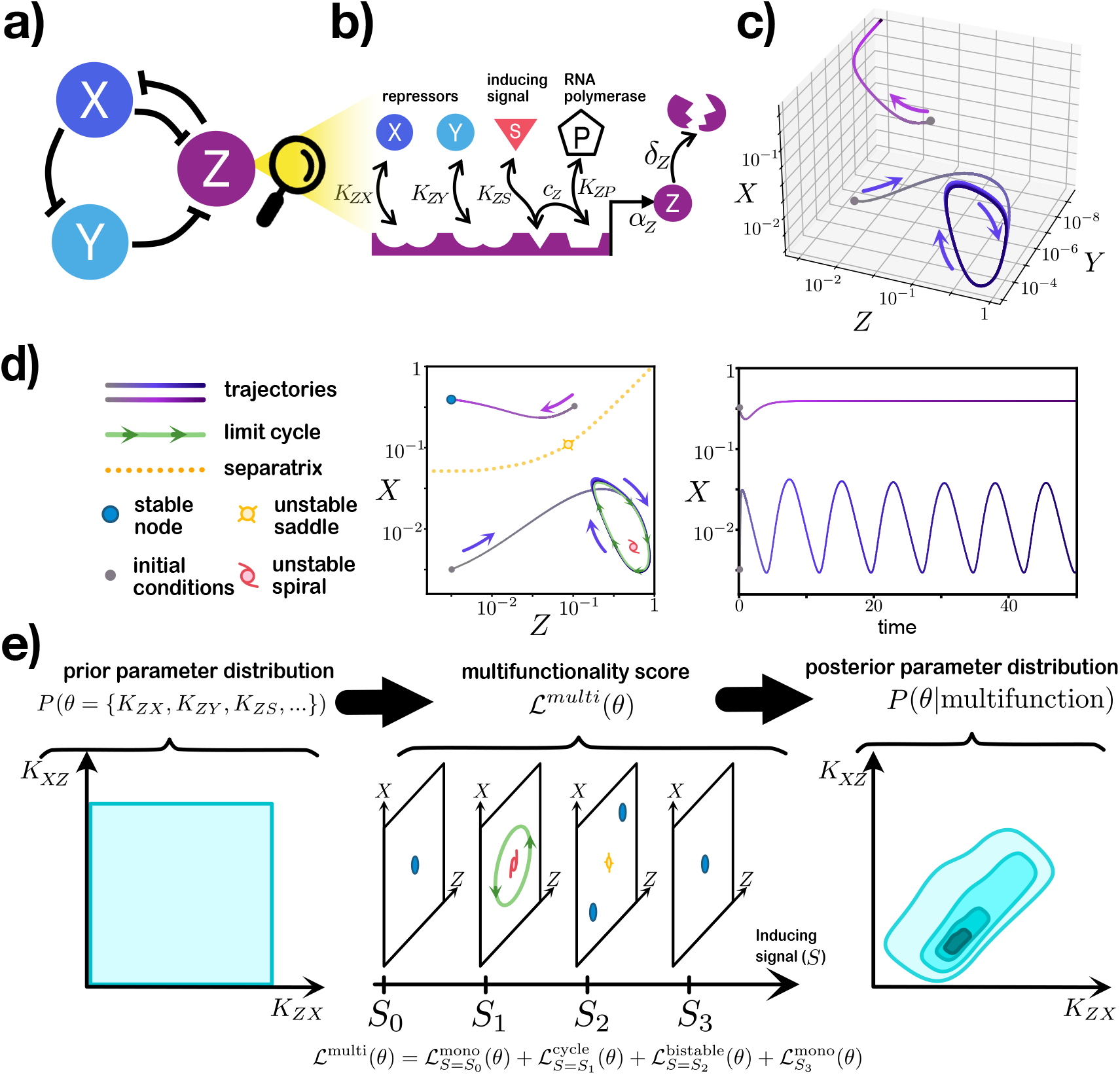
The AC-DC is a multifunctional gene regulatory motif. (a): Topology of the AC-DC circuit. (b): Gene expression is modelled using a thermodynamic formalism that enumerates all possible promoter-bound states and their corresponding equilibrium energies. (c): Example trajectories in a parameter regime where the circuit exhibits bistability between oscillations (blue) and a stable steady state (purple). The resulting behaviour depends on the system’s initial conditions (grey circles). (d): Dynamical analysis of the AC-DC behaviour in panel c, showing equilibria including steady states (stable and unstable) and limit cycles. (e): Parameter space encoding multifunctionality can be uncovered using Approximate Bayesian Computation, transforming a flat parameter distribution (uninformative prior) into a posterior distribution that is compatible with a multifunctionality score, resulting from imposing dynamical behaviours at different values of the external signal. All the parameters used in this Figure are summarized in supplementary table II for *S* = 800.

Despite its theoretical promise, and a synthetic implementation of the topology with some multifunctional features [28], its full dynamical potential remains largely unexplored. The inherent fragility of multifunctional circuits – where small parameter changes can completely alter behaviour – poses a substantial challenge. This sensitivity, demonstrated experimentally through strong epistatic effects even in small networks [29], complicates efforts to predict how combinations of mutations or design choices will impact function. These observations highlight the importance of developing mathematical models that use realistic regulatory functions when modelling GRNs. For example, the choice of sigmoidal response (such as the Hill function) can substantially influence model outcomes. To address this, thermodynamic models [30–33] or non-specific binding equilibrium models [34, 35] have been proposed as simple yet biophysically grounded formalisms that capture biological nonlinearities without adding mathematical complexity.

In this work, we take a systematic and exploratory approach to uncover the hidden dynamical landscape of the AC-DC circuit. We employ a thermodynamic formalism to describe regulatory interactions more realistically than classical Hill-based models and focus on simplified inducer architectures that reduce experimental complexity, paving the way for future experimental implementation. By leveraging Approximate Bayesian Computation, we rigorously map the parameter spaces supporting diverse dynamic behaviours and assess their robustness. Surprisingly, we find that using simpler and more biophysically accurate parameters preserves the AC-DC dynamical behaviours. In addition, we find that the simpler topology is able to expand upon the known range of attainable behaviours, thereby revealing an unexpected richness of multifunctional potential for this topology. Our results provide concrete design guidelines for building AC-DC circuits in the laboratory and open new avenues for both synthetic circuit engineering and the understanding of emergent dynamics in natural gene regulatory systems.

## II. RESULTS

### A. Biophysical exploration

The dynamics of the AC-DC circuit can be described using ordinary differential equations (ODEs) that represent the expression levels of each of the three nodes, X, Y, and Z, under the influence of an external inducing signal *S*:

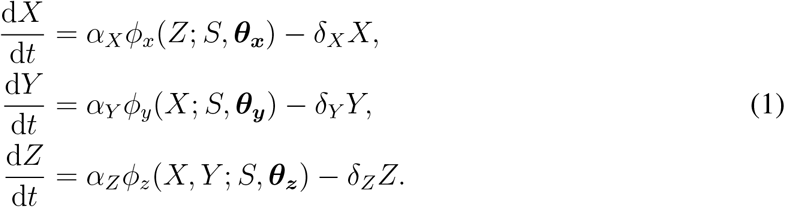

Where parameters *α* and *δ* correspond with the maximal production and degradation rate for each transcription factor respectively (see Fig. 1b). The regulatory logic is encoded in the functional form of the regulatory function *ϕ*(·) and the parameter vector ***θ***. While previous models have used Hill functions [36] or generalized linear models [27], we employ a thermodynamical description (see Fig. 1 b and Appendix A 1) based on enumerating all the possible promoter occupancy states and their binding energies [37–39] that allow us to calculate the probability of finding the gene in an actively transcribing state (bound to an RNA polymerase). In this formalism, each parameter in ***θ***_***j***_ has a biophysical interpretation, and the non-linearities arise naturally from the prescribed molecular interactions without the need to impose Hill exponents. Specifically, each gene is characterized by a set of binding affinities *K*_*ji*_ of molecule *i* to promoter *j*; for example *K*_*XP*_ denotes the binding affinity of polymerase to the empty promoter of gene X, and *K*_*ZY*_ is the binding affinity of protein Y to promoter Z. In this formulation, the role of the inducing signal *S* is reducing the binding energy of RNA polymerase through the cooperativity parameter *c*_*i*_.

Explicit forms of the function are provided in the Supplemental Material section A, which includes 13 distinct parameters for the two-inducer circuit (17 before non-dimensionalisation).

An initial exploration of the model confirms its ability to reproduce the bistability between oscillations and a steady state, consistent with the dynamical behaviour reported previously [36] (see Fig. 1 c-d). However, uncovering the full range of behaviours accessible to this formulation requires a more systematic exploration of its high-dimensional parameter space. To this end, we employ a Bayesian inference strategy using a sequential Monte Carlo scheme, which allows us to progressively chisel out regions of parameter space compatible with prescribed dynamical behaviours. In particular, we define a score function that quantifies the distance to a target multifunctional behaviour given a set of parameters. For example, in the first part of this work, the multifunctional AC-DC behaviour is defined through a set of dynamical regimes for different values of the inducer signal *S* – namely a combination of monostable, bistable, and oscillatory regimes (see Fig. 1e). Explicit forms of the score function are provided in Appendix A 3. One key advantage of this approach is that it does not attempt to find a single point estimate of a parameter set that fulfils the dynamics but instead identifies regions of parameter space that support a prescribed multifunctionality. This enables us to assess the robustness of each behaviour, facilitates model selection, and ultimately provides concrete design guidelines.

### B. One-inducer vs two-inducer

AC-DC dynamics have been captured in the literature using two-inducer models where signal activation acts on two genes of the circuit (see Figure 2a), inspired by the original network topology guiding neural progenitor differentiation [26]. The question remains whether there is a simpler system capable of exhibiting the same multifunctional dynamics. For this, we consider a one-inducer system (see Fig. 2a) where signal activation acts only on gene X. Using the multifunctional Bayesian exploration (see Appendix A 3) we find that multifunctional dynamics are exhibited by both oneand two-inducer systems. We expect that two-inducer systems can be achieved in broader regions of the parameter space, since the one-inducer system is a subcase of the two-inducer system. This was confirmed when examining marginal and pairwise posterior distributions for common parameters in both models (see Fig. 2b and Fig. A5).

**FIG. 2.**
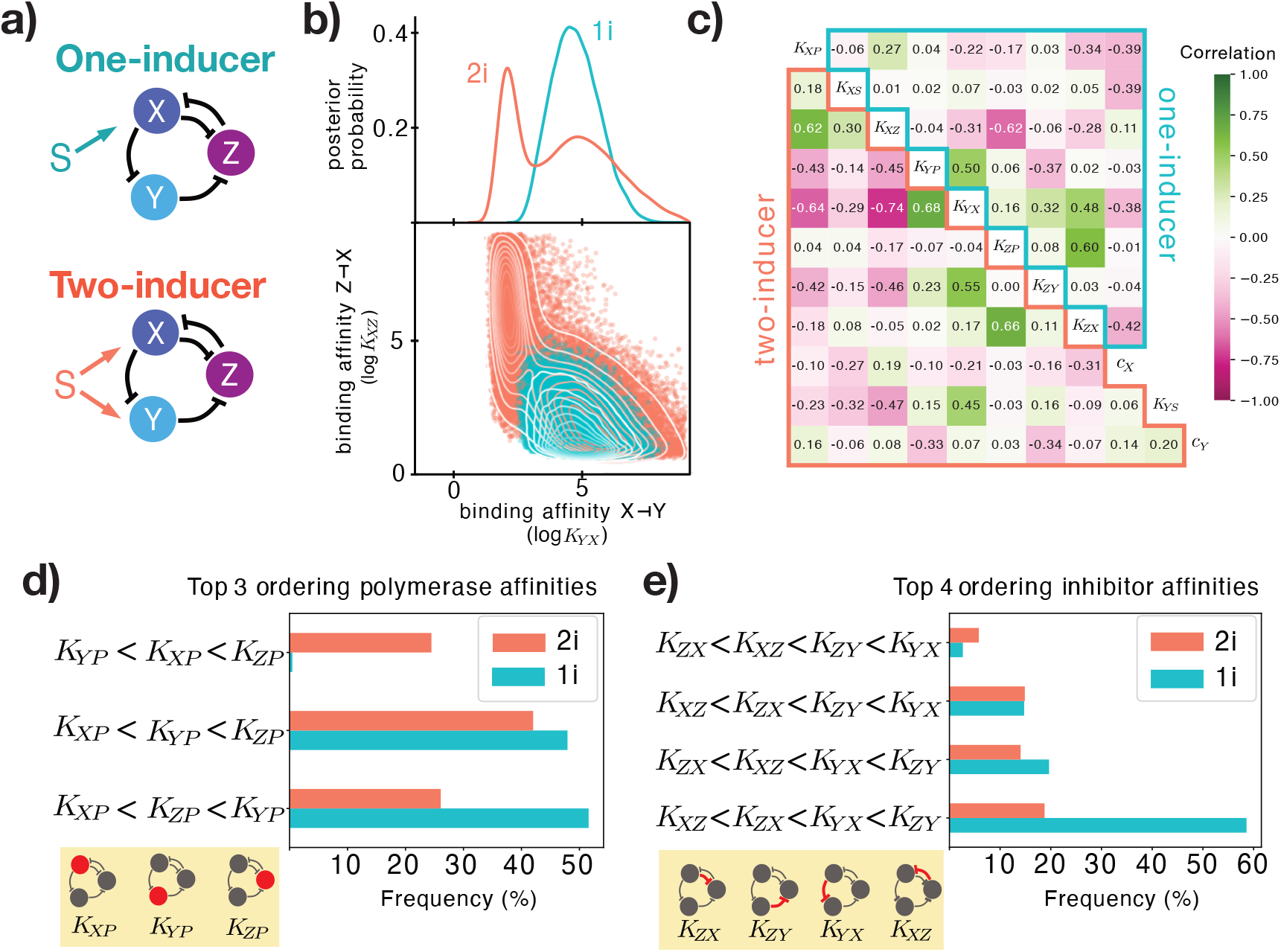
(a): One- and two-inducer AC-DC circuits. (b): Posterior probability distributions comparing one-inducer (1i, blue) vs two-inducer systems (2i, pink), including marginal posterior distribution for the inhibitor binding *K*_*Y X*_ (top) and bivariate distribution for *K*_*Y X*_ and *K*_*XZ*_ (bottom). (c): Correlation of the posterior distribution of each pair of parameters for the one-inducer (top-right) and two-inducer systems (bottom-left). (d): Top three relative polymerase binding intensity orderings. (e): Top four relative inhibitor binding intensity orderings.

However, the 2-inducer model includes two additional parameters, *K*_*Y S*_ and *c*_*Y*_, which makes its parameter space higher-dimensional. As a result, a broader spread across the shared dimensions of the respective hypervolumes does not, by itself, reveal which circuit is more prone to multifunctional behaviour – that is, in which circuit multifunctionality is more likely to appear when parameter sets are sampled at random. To address this, we compared the relative hypervolumes of the posterior distribution with respect to their prior (analogous to a Bayes factor; see Appendix A 4 for details). This analysis showed that the one-inducer model is 1.5 times more likely to show multifunctionality than a two-inducer model. This implies that once dimensionality is taken into account, the one-inducer model has a slightly larger effective spread than the two-inducer model. This conclusion is further supported by the pairwise posterior parameter correlations (Fig. 2c), where the two-inducer model displays stronger correlations. High correlations, such as between *K*_*Y X*_ and *K*_*XZ*_, indicate fragility to parameter perturbations, requiring coordinated tuning of multiple parameters to make the two-inducer circuit viable for synthetic development.

Given the wide parameter ranges compatible with multifunctionality, we explored the parameter space to uncover structural features associated with this behaviour. This analysis revealed that the relative ordering of promoter strengths (polymerase binding rates *K*_*XP*_, *K*_*Y P*_, *K*_*ZP*_) plays a key role, with only three dominant orderings observed, and just two in the one-inducer system corresponding with *K*_*XP*_ being the weakest promoter (see Fig. 2d). Similarly, analysis of repressor binding strengths revealed four dominant inhibitor binding orderings (Fig. 2e). While two-inducer systems tended to favour similar inhibitor orderings, the one-inducer system showed a clear preference for the configuration *K*_*XZ*_ *< K*_*ZX*_ *< K*_*Y X*_ *< K*_*ZY*_, suggesting specific rules for constructing a functional one-inducer circuit. Since the simpler one-inducer circuit already displays clear multifunctional behaviour, we will focus the rest of our analysis on this simpler circuit unless otherwise noted.

### C. One multifunctionality is compatible with different criticality behaviours

The definition of multifunctionality used in our parameter search focused on prescribing specific dynamical behaviours at defined levels of the inducing signal (e.g. number of steady states or existence of a limit cycle, see Fig.1e). Crucially, this definition is agnostic to the specific bifurcations through which these behaviours emerge, i.e. it does not constrain the transitions by which the number or nature of attractors changes with the inducing signal.

To investigate this, we focused on the transition from limit cycle oscillations to bistability as the inducing signal increases from *S*_1_ to *S*_2_ (depicted in Fig. 3a, cf. 1e). As expected, bistability emerges from a saddle-node bifurcation. However, oscillations can disappear via distinct bifurcation patterns [8]. We observe three different transitions out of the oscillatory regime (Fig. 3b): (i) a Saddle-Node on Invariant Circle bifurcation (SNIC), where the saddle-node bifurcation splits the limit cycle; (ii) a saddle-loop bifurcation, in which the cycle loses stability by colliding with the saddle point introduced by the saddle-node bifurcation; and (iii) a supercritical Hopf bifurcation, where the limit cycle gradually shrinks and vanishes.

**FIG. 3.**
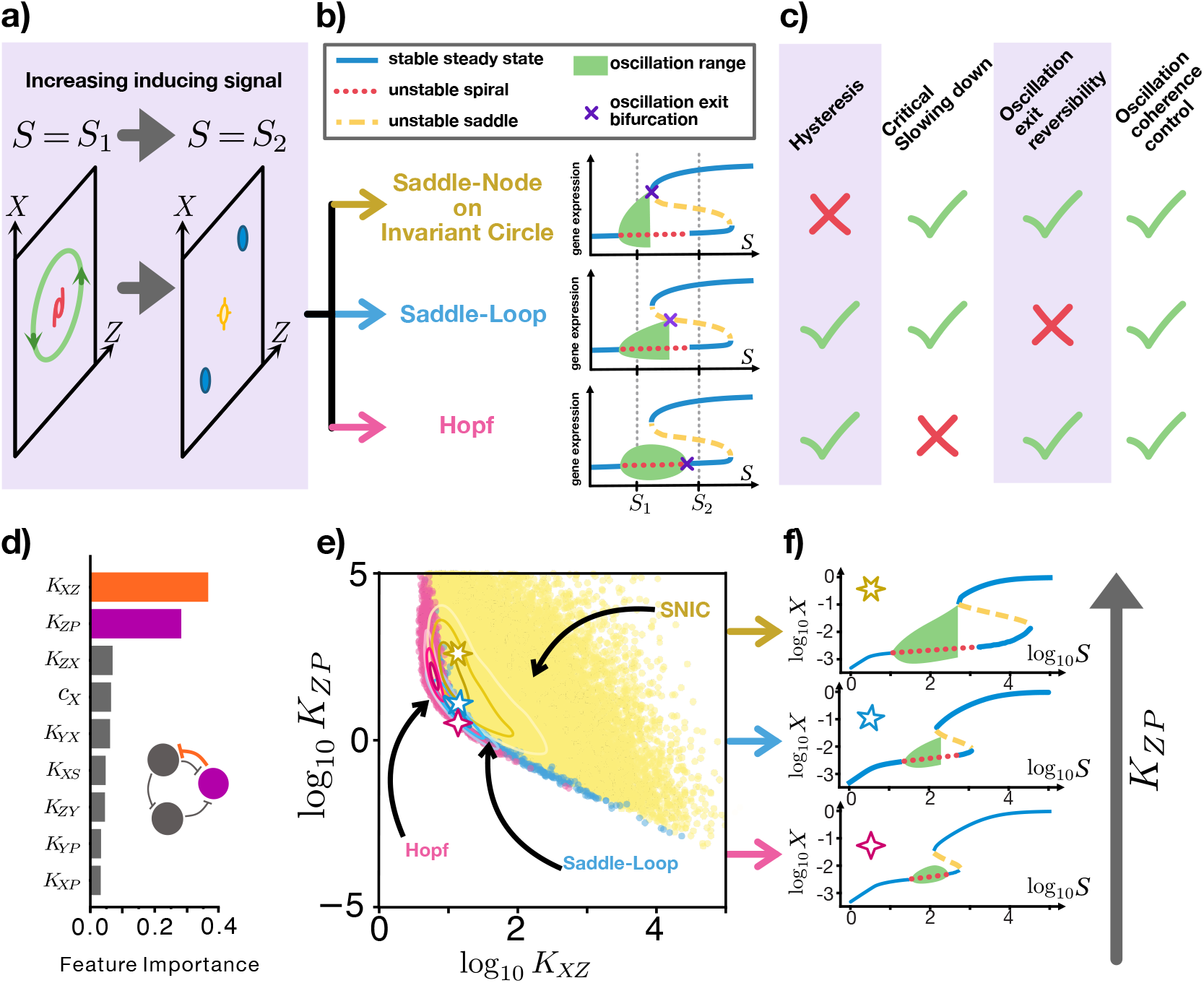
The AC-DC transition from oscillations to bistability can show 3 different critical behaviours. (a): The condition for multifunctionality requires that at two different signal levels, the system goes from stable oscillations to a bistable behaviour. Line and marker meaning is the same as in Fig. 1d. (b): The transition from oscillation to bistability can be classified by three different bifurcation patterns in which oscillations (green area) disappear as the signal is increased continuously. Each oscillation exit-point as signal is increased is marked with a purple cross. (c): Each bifurcation diagram has different dynamical properties despite having the same multifunctional prescription. (d): Decision tree classification of the three bifurcation categories identifies *K*_*XZ*_ and *K*_*ZP*_ as the main parameters in charge of controlling the resulting bifurcation diagram. (e): Scatter plot for the pairwise posterior joint distribution of parameter pairs (*K*_*XZ*_, *K*_*ZP*_), coloured by the resulting bifurcation diagram. Lines indicate the high density regions corresponding to each diagram. Stars mark the parameter sets used in panel f. (f): The three bifurcation diagrams can be obtained by changing *K*_*ZP*_. The shown diagrams correspond to log_10_ *K*_*ZP*_ = [2, 0.7, 0.5]. All further parameters can be found in Table II.

Far from a purely mathematical exercise, different bifurcations give rise to distinct dynamical behaviours (Fig. 3c). For instance, the SNIC bifurcation is the only case in which the limit cycle disappears precisely at the signal value where a new steady state appears, eliminating the possibility of bistability and hysteresis between the oscillatory and steady-state regimes. On the other hand, both the SNIC and the saddle-loop bifurcations are homoclinic and introduce critical slowing down near the bifurcation point, where the period of oscillations diverges (this critical slowing down is further studied in Section II D) [40]. This is not the case for the Hopf bifurcation, in which the period remains constant as the amplitude gradually shrinks. Another important difference concerns the reversibility of the transition out of oscillations. In both the SNIC and Hopf cases, oscillations can re-emerge with only a small reduction of the signal. In contrast, after a saddle-loop bifurcation the system settles into a different steady state, and oscillations can only be recovered through a much larger change in signal.

Note that all these dynamical behaviours are compatible with other idiosyncratic behaviours already reported for the AC-DC, such as the control of oscillation coherence [36]. In addition, note that in this classification we are not interested in the bifurcations by which the unstable spiral changes stability (e.g. change in stability from unstable to a stable steady state at higher values of *S* in the three diagrams of Fig. 3b), since the new stable branch is not accessible through continuous changes in the inducing signal. Bifurcations around this inaccessible point, can include an additional Hopf bifurcation, introducing further regimes of bistability between a limit cycle and a steady state. These have been omitted in the bifurcation diagrams for the sake of clarity.

Interested by this variability of behaviours we proceeded to analyse which parameters are in charge of producing each specific bifurcation case. Using a decision tree classifier we identified that two parameters are mostly involved in controlling this behaviour. In particular the strength of the promoter Z (*K*_*ZP*_) and the repression strength of gene Z (*K*_*XZ*_) (see Fig. 3d).

Further analysis of the three categories across the successful parameter sets reveals that the SNIC bifurcation is the most common scenario (Fig. 3e and Fig. A3). Similar results were obtained in the 2-inducer case, confirming that both the 1-inducer and 2-inducer systems have comparable dynamical capabilities (see Fig. A3). Moreover, our analysis shows that the three behaviours occupy neighbouring regions in parameter space (Fig. 3e), suggesting that transitions between them can occur continuously with minimal parameter changes, a signature of tuneability of the dynamic features of the circuit. This was confirmed by continuously increasing the value of *K*_*ZP*_ in a system initially exhibiting the Hopf scenario, which sequentially transitioned through a saddle-loop and ultimately into a SNIC regime (see Fig. 3f).

### D. Choice of bifurcation allows for the control of critical slowing down properties

Although the three different bifurcation categories highlighted in the previous section may share the presence of certain dynamical features (Fig. 3c), the specific characteristics of these features can differ substantially across categories. For instance both the saddle-loop and the Hopf exit allow for bistability between oscillations and a steady states, nevertheless, we expect the Hopf exit to have larger signal ranges of bistability. These differences reveal the importance of choosing specific bifurcation diagrams where the robustness of a specific dynamical behaviour is required.

A case of specific relevance is the critical slowing down observed in both SNIC and saddle-loop bifurcations. In these cases, the bifurcation occurs via the formation of a homoclinic orbit (Fig. 4a), leading to a divergence of the oscillation period as the signal approaches the critical value *S*_crit_ (Fig. 4b). Notably, in the saddle-loop case, the trajectory passes near a saddle point even before the bifurcation occurs, potentially slowing the cycle across a broader range of signal values (Fig. 4a). To test this, we analysed all parameter sets exhibiting each bifurcation and measured how the oscillation period changes with the signal near *S*_crit_, quantified as the critical slope *m* (Fig. 4c). The results confirm that saddle-loop bifurcations consistently display smaller values of *m*, indicating a larger signal range required for the variation of the period near the transition point (Fig. 4d). By contrast, SNIC bifurcations show a steeper slope, suggesting a more stark dependence with the signal, and therefore less dynamic control over the period. These results highlight the saddle-node as a favourable mechanism for modulation of oscillatory periods in gene circuits.

**FIG. 4.**
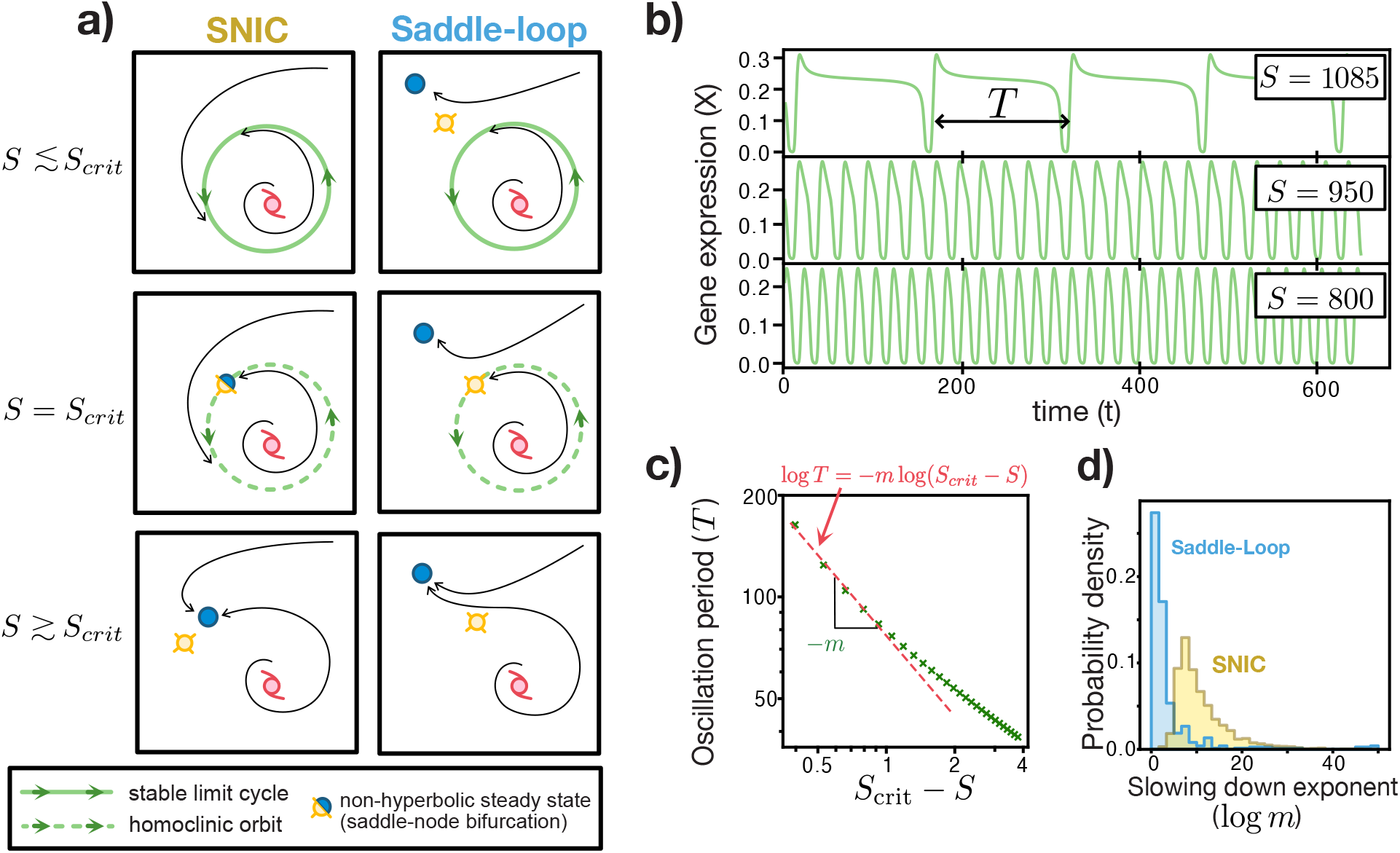
Slowing down properties near SNIC and saddle-loop bifurcations. (a): Phase portraits of the SNIC and Saddle-loop bifurcations as they cross the bifurcation signal *S*_*crit*_. Black arrows indicate the flow of the system for different initial conditions. All the symbols not indicated in the legend correspond with the ones in Fig. 1d. (b): Trajectories of expression of gene X in time, showing the slowing down of the oscillations as the the signal approaches *S*_*crit*_ = 1086.95. (c): Change of the oscillation period with the distance to *S*_crit_. Red dotted line shows the log-log linear fitting use to calculate the slowing down exponent *m*. For small *m* the slow down occurs through a larger range of *S*, while for large *m* there is comparatively significant slowing down with a small change in the signal. (d): Probability distribution of slopes *m* for both bifurcations. Parameters for b and c are given in Table II.

### E. A Zoo of multicriticalities

In the previous sections, we examined how sustained oscillations transitioning into bistability can arise through different bifurcation routes, each with distinct dynamical features. Building on this, we sought to explore more broadly the types of bifurcation diagrams compatible with the AC-DC circuit. To do so, we relaxed the multifunctional constraints defined in Fig.1e, and instead searched for any solution satisfying three general multifunctional conditions: (1) monostability at low and high signal levels, (2) the existence of a stable oscillatory regime (i.e., a limit cycle), and (3) bistability between two attractors. These criteria do not enforce a fixed sequence between oscillations and bistability or restrict bistability to steady states alone (see Appendix A 3).

Following the same strategy as in the previous sections, we classify the resulting dynamical behaviours based on how oscillations emerge and disappear as the inducing signal increases continuously from the monostable regime at low signal to the monostable regime at high signal. The possible ways in which oscillations can end are the same as those discussed in Section II C, namely, via a Hopf bifurcation, a SNIC, or a saddle-loop bifurcation. Likewise, oscillations can emerge through three distinct mechanisms: a Hopf bifurcation, a SNIC, or a saddle-node bifurcation that induces a jump to a pre-existing limit cycle (Fig. 5 a), the latter being a type of bifurcation that initiates oscillations but cannot terminate them.

**FIG. 5.**
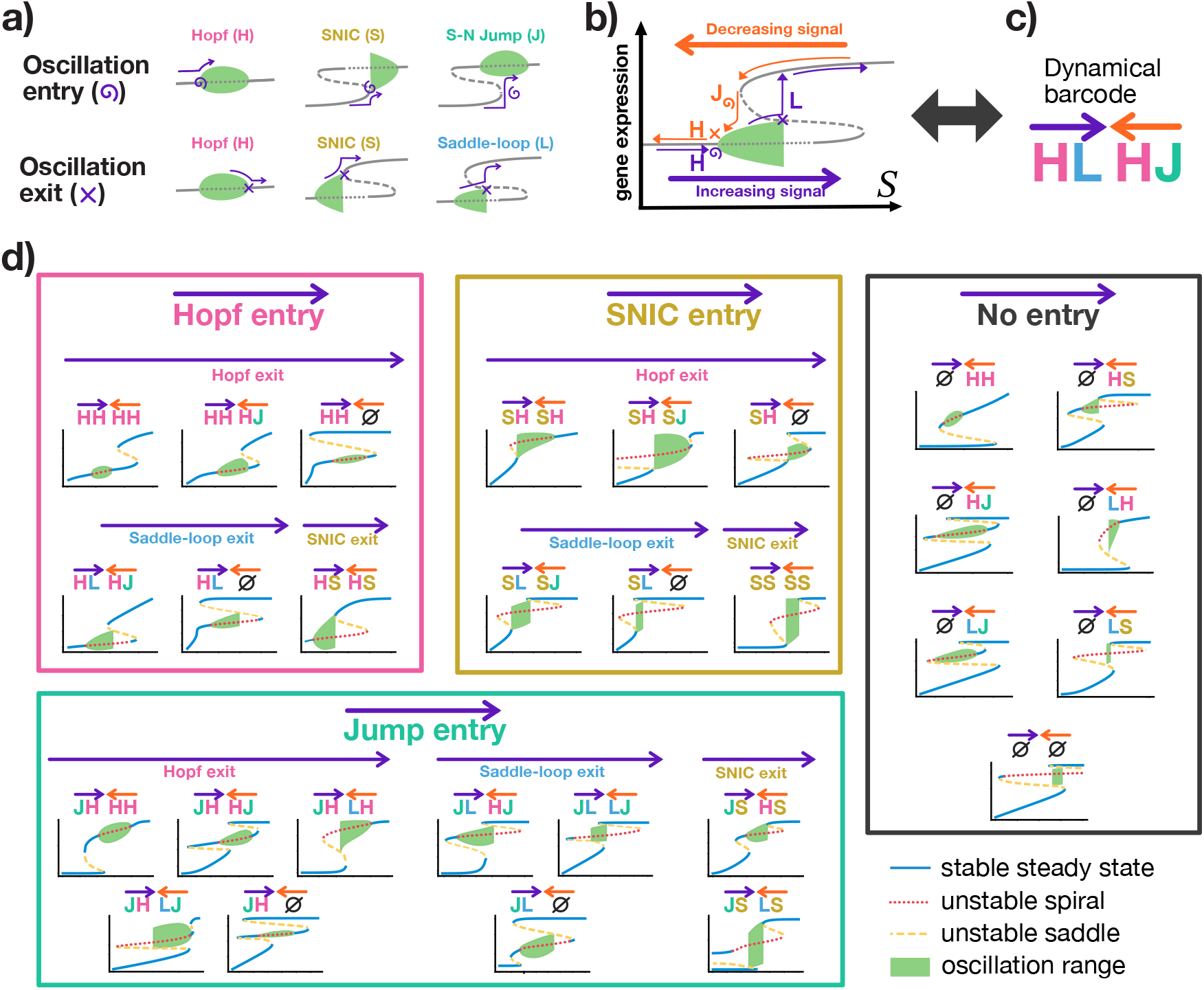
The AC-DC circuit is compatible with 29 different multifunctional behaviours. (a): Via a continuous modulation of the inducing signal oscillations can start or end through three different bifurcation events each. (b): Schematic showing that the bifurcations leading to the start and end of an oscillatory behaviour depends on the history of the inducing signal. (c): All the possible dynamical behaviours can be encoded in a dynamical barcode that captures how oscillations start and end as signal is increased from low to high levels, or decreased from high to low levels. (d): Resulting bifurcation diagrams (gene expression *X* against signal *S*) from the parameter search for each one of the 29 dynamical barcodes found. Parameters and ranges for each diagram are available in the provided code repository.

Interestingly, due to the hysteresis introduced by certain bifurcations, the progression of oscillatory behaviours differs depending on whether the signal increases from low to high or decreases from high to low. This can be seen in Fig. 5b, where, by increasing the signal, oscillations start with a Hopf bifurcation and end with a saddle-loop bifurcation. On the other hand, as signal is reduced back, oscillations start with a saddle-node jump into a cycle, and end with a Hopf bifurcation. This has important implications for multifunctionality, as it allows the same circuit to access distinct dynamical properties depending on the direction of signal change. To capture this, we label the dynamical repertoire of each bifurcation diagram by enumerating the four bifurcations through which oscillations start or end as the signal increases or decreases. This results in a “dynamical barcode” that uniquely characterises the oscillatory regimes accessible to each diagram (Fig. 5c).

Surprisingly, the result from this classification returned 29 different dynamical barcodes observed in our parameter search (Fig. 5d). Interestingly some of the behaviours involve scenarios in which the oscillations are not accessible in specific directions of the signal e.g. no oscillatory regime is observed when increasing the signal from low to high values, but oscillations appear when the signal is reduced from high to low values (e.g.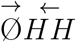). The system also shows the possibility of tristability between oscillations and two steady states (e.g. 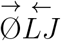), expanding the known behaviours from the traditional bistable switch component of the AC-DC.

## III. DISCUSSION

In this study, we explored the rich dynamical repertoire of the AC-DC circuit using a biophysical formulation that brings greater mechanistic realism compared to previous approaches. Remarkably, we show that the circuit not only retains the multifunctionality previously reported, but that these behaviours also emerge in a simpler version of the circuit requiring only a single signal input, rather than relying on multiple inducers [5, 36] or embedding in a larger network [27]. This has important implications both for understanding the role of AC-DC-like motifs in natural developmental circuits [27, 41] and for synthetic biology, where building robust multifunctional circuits from minimal components is a key design goal [42–44]. Moreover, our use of Approximate Bayesian Computation offers a practical route toward rational circuit design, returning explicit ranges and orderings of binding affinities and promoter strengths that support targeted dynamical behaviours. This approach makes the circuit more accessible to implementation using standard synthetic biology toolkits, facilitating design cycles that combine model-driven inference with measurable biophysical parameters [45] and overcome model non-identifiability common in ODE driven predictions in systems biology [46].

Our results also highlight how the choice of regulatory functions can strongly influence mechanistic predictions. For example, while previous studies identified the saddle-loop bifurcation as the key route connecting oscillations and bistability [7], in our model the SNIC bifurcation emerged as the most plausible transition. This aligns with recent work in embryonic development, where SNIC transitions have been proposed to regulate oscillations in the segmentation clock [47, 48], heart beat initiation [49] and stem cell differentiation [8]. However, the prevalence of SNICs in our study may be driven by symmetric assumptions, such as equal degradation rates across all proteins, a common simplification known to stabilise oscillations [50]. In contrast, our extended parameter scans required heterogeneous degradation rates to uncover the full bifurcation zoo, again demonstrating how structural assumptions in ODE models can shape the biological interpretations. Given the different functional consequences of SNIC and saddle-loop bifurcations, a future systematic exploration of these structural determinants will be critical.

One of the most unexpected findings of our study is the sheer diversity of accessible dynamical behaviours. The 29 bifurcation diagrams we report represent only a subset of the AC-DC circuit’s full potential. For instance, we excluded simpler dynamics such as globally monostable systems, bistable systems that are non-oscillatory, or oscillatory systems without bistability (Fig. A2a-c), as well as diagrams with bistability in the absence or saturation of signal, which could enable irreversible transitions (Fig. A2d), a behaviour observed in theoretical and experimental studies of bistable switches in synthetic [23] and stem cell biology [51]. Including these would expand the set of circuits far beyond 29. Notably, under our constraints, 31 dynamical barcodes are theoretically feasible. The two missing barcodes (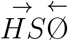 and 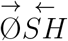 presented in Fig. A2e) were not observed in our search, possibly due to parameter sampling limitations or because they are structurally inaccessible, requiring future analytical work.

Some circuits displayed particularly intriguing behaviours. For instance, diagram with barcode 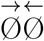 shows an oscillatory regime that is not accessible via changes in the inducing signal. This behaviour is reminiscent of isola-type dynamics recently described in mushroom bifurcations [24, 52]. In such scenarios, oscillations can only be reached by tuning a secondary parameter that allows transitions between bifurcation diagrams, effectively locking the system into an oscillatory state that can be irreversibly destroyed by modulating the main signal. Similar behaviours are found in several of the 29 diagrams, suggesting that the AC-DC circuit can be tuned to function as a timer, memory element, or both. Recent research further shows that combinations of nearby bifurcations can act synergistically to prolong transients over broad signal ranges, offering finer control of oscillatory periods than the mechanisms discussed in Section II D [25]. Similarly, future work could explore how environmental fluctuations or stochastic gene expression affect these bifurcation patterns, and whether such effects can be harnessed for probabilistic decision-making in synthetic systems.

More broadly, this work underscores the functional complexity embedded even in small regulatory motifs, a fact that should not be disregarded when reverse-engineering networks from high-throughput data, and encourage careful consideration of regulatory functions when drawing mechanistic conclusions or predictions out of inferred networks.

## ACKNOWLEDGMENTS

R.P.-C., S.M., and A.P. acknowledge funding by the Leverhulme Trust Research Project Grant No. RPG-2023-085. Y.S. and I.A. acknowledge the Swiss National Science Foundation (320030-236359 awarded to Y.S.), and M.M. Fischer gratefully acknowledges funding by the Deutsche Forschungsgemeinschaft (DFG, RTG2424 “CompCancer”, project number 377984878).

## Appendix A: Methods

### 1. Thermodynamic formalism

To simulate gene regulatory dynamics, we employ a thermodynamic formalism [37–39, 53], also known as a fractional occupancy model, that describes the binding events of DNA, transcription factors (TF), and RNA polymerase in biophysical terms. In this framework, gene expression corresponds to the probability that the promoter is in a state with a bound RNA polymerase. This involves enumerating all possible promoter occupancy states and calculating the fraction that include a bound polymerase (Fig. A1). The relative abundance of each state depends on its binding energy, with transcription factors and signals modulating the energy of different configurations.

The outcome is a transcription probability function for each promoter (or node) in the system. For example, node *i* is characterised by a probability *ϕ*_*i*_(*X, Y, Z, S*) that depends on the binding affinities *K*_*ij*_ of molecule *j* to promoter *i*, as well as a cooperativity parameter *c*_*i*_ describing the energetic facilitation of polymerase binding once the inducer signal is bound. For instance for the 1-inducer model (Fig. A1) the probability is described as

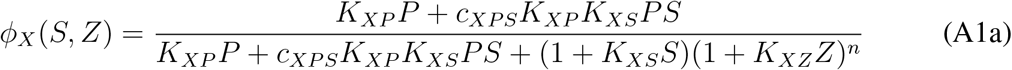

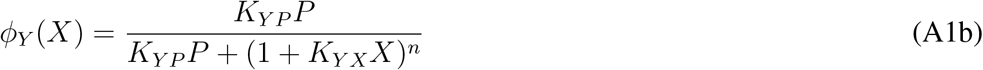

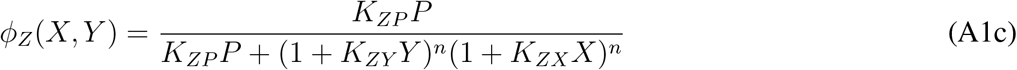

In this formulation, the repressive TF forbids the binding of polymerase, where *n* is the number of binding sites. For the simulations in this manuscript we fixed *n* = 3. In addition, since the total polymerase *P* is kept constant in our simulations, we set *P* = 1 without loss of generality, measuring the binding affinity *K*_*iP*_ in units of polymerase concentration. Similarly, for the 2inducer case, the only change is on the extra signal binding to promoter Y,

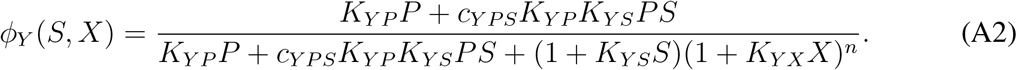

The resulting non-linear transcription probabilities *ϕ*_*i*_ can be directly incorporated into the ODEs (Eq. 1), allowing us to simulate the system’s dynamics. Additionally, for our parameter search we normalise the expression levels by their maximum value, using the dimensionless levels

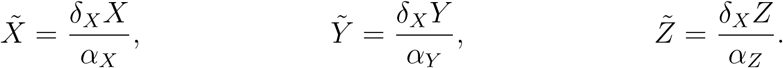

**FIG. A1.**
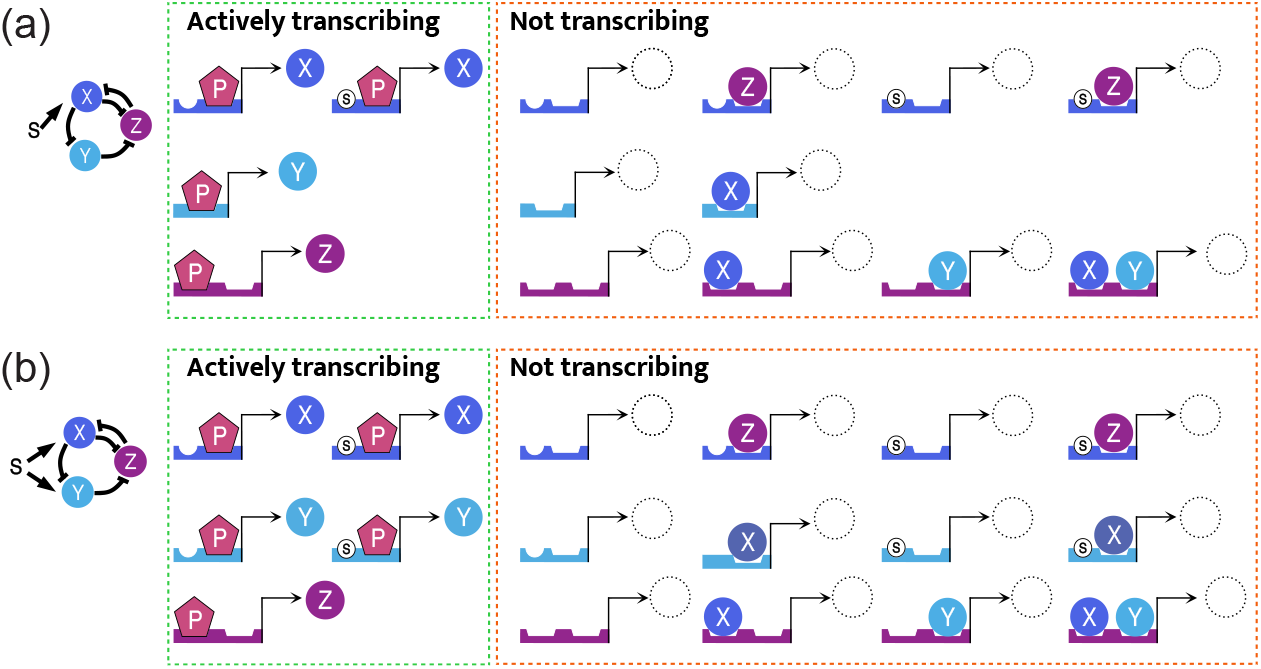
Thermodynamical formalism configurations for the AC-DC models. For all three nodes, schematics of all possible configurations of promoter binding states are displayed, indicating whether they are actively transcribing (RNA polymerase bound) for (a) the one-inducer model and (b) the two-inducer model.

Furthermore, we reparametrize time 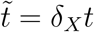 resulting in dimensionless degradation rates 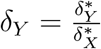 and 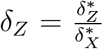. For the sake of clarity, we will drop the tildes for all the dimensionless parameters ad variables. These reparametrisations reduce the number of free parameters from 17 (15 in the one-inducer model) to 13 (11 in the one-inducer model).

### 2. Bifurcation analysis

In order to obtain the bifurcation diagrams, we find the steady states (*X*^*^, *Y* ^*^, *Z*^*^) and corresponding linear stability of dimensionless version of the system (eq. 1) at different values of *S*. The topology of the AC-DC allows us to find the steady states by numerically finding the solutions of the the 1-dimensional equation

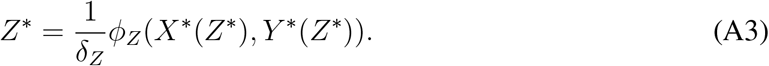

Here *X*^*^(*Z*^*^) and *Y* ^*^(*Z*^*^) correspond with

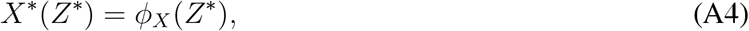

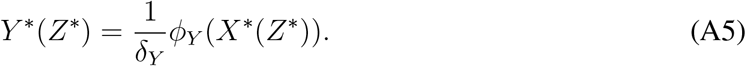

Limit cycles are obtained by calculating long-time trajectories originating from perturbed unstable points. Bifurcation points are characterised as points where the stability of a branch changes (e.g. Hopf bifurcation), points where the number of steady states changes (e.g. saddle-node bifurcation), or points at which a limit cycle becomes a homoclinic orbit (e.g. saddle-loop bifurcation, saddle-node on invariant circle)

### 3. Parameter exploration

We explore parameter space using Approximate Bayesian Computation using a sequential Monte Carlo-based sampling approach [54, 55], using flat prior distributions along the parameter ranges indicated in table I. Distance functions *d* (inverse of score functions / pseudo-likelihood ℒ) are introduced as a metric for the sample rejection based on the dynamics obtained from steady state analysis. The distance functions determine how far a parameter set is from the desired dynamical properties, with larger values indicating that the desired dynamical criteria are not being met. We evaluate the phase portraits at *S* = 1, *S* = 10^2^, *S* = 10^3^, *S* = 10^4^ (see Fig. 1e) to obtain a multifunctionality score.

The oscillatory behaviour is evaluated at *S* = 10^2^, imposing the presence of an unstable spiral. Multiple equilibria are penalised, as well as monostability in the absence of a complex component to the eigenvalues. We also prioritise eigenvalues *λ* = *a* ± *iω*, that result in faster repulsion and higher oscillation frequency to select for biologically observable oscillations.

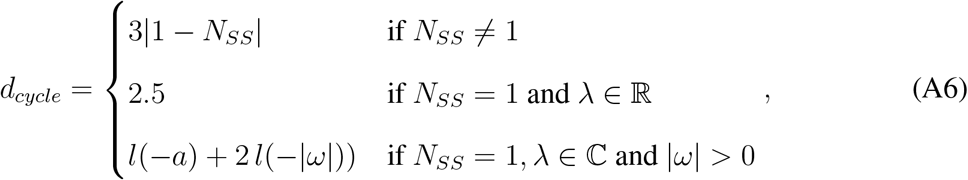

where *N*_*SS*_ is the number of steady states, and 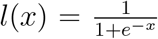 is the logistic function to normalize the distance metric. Similarly, monostability behaviour at *S* = 1 and *S* = 10^4^ is selected by penalizing multistability and favouring non-oscillatory, fast nodes.

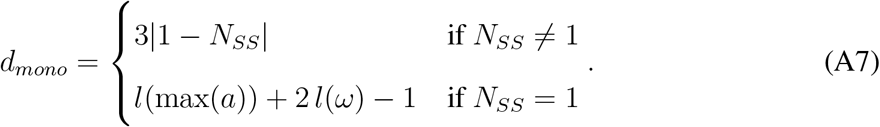

We also evaluate for bistability at *S* = 10^3^, selecting for the coexistence of three steady states of undefined nature.

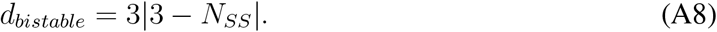

The final multifunctionality score for each parameter set corresponds to the sum

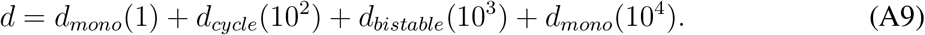

For the one-inducer versus two-inducer analysis in section II B we generate 4 × 10^4^ parameter sets (per circuit) which satisfy a final distance function threshold *d* = 1.35.

For the results presented in the bifurcation zoo (Fig. 5) our aim was not estimating posterior distributions, but uncovering the existence of specific bifurcation diagrams. To this end, we extended the ABC search using a semi-automatic approach. We first broadened the initial ABC screening by modifying the distance function allowing for other combinations of multifunctionality, and then performed targeted manual exploration around specific parameter sets to sharpen the observed behaviours or to reveal nearby alternatives. For example, varying degradation rates induces transitions between related dynamical barcodes related to oscillation amplitude. Likewise, modulating signal–polymerase cooperativity significantly alters the ranges of multistability. Other parameter variations allow fine-tuning of individual branches and shifting the signal ranges over which multifunctionality emerges.

Initial parameter sets leading to 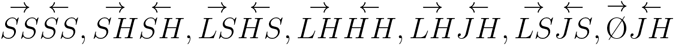 behaviours were obtained with the following inverted distance function.

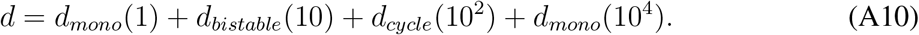

For initial 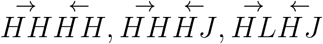, and 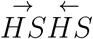, we used smaller values of input signal using the distance function

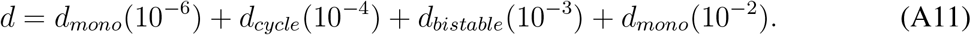

This classification for the search does not include trivial cases that are also compatible with the AC-DC topology such as monostability, oscillations without bistability, or multistability without oscillations (see Fig. A2a-c). Similarly, the search ignores cases in which the system is not reset to monostability at high and low signals resulting in irreversible scenarios (see Fig. A2d). Finally, there are two bifurcation diagrams that even though they fulfill the conditions of the zoo (with the corresponding barcode), they were not found in the search (see Fig. A2e).

**FIG. A2.**
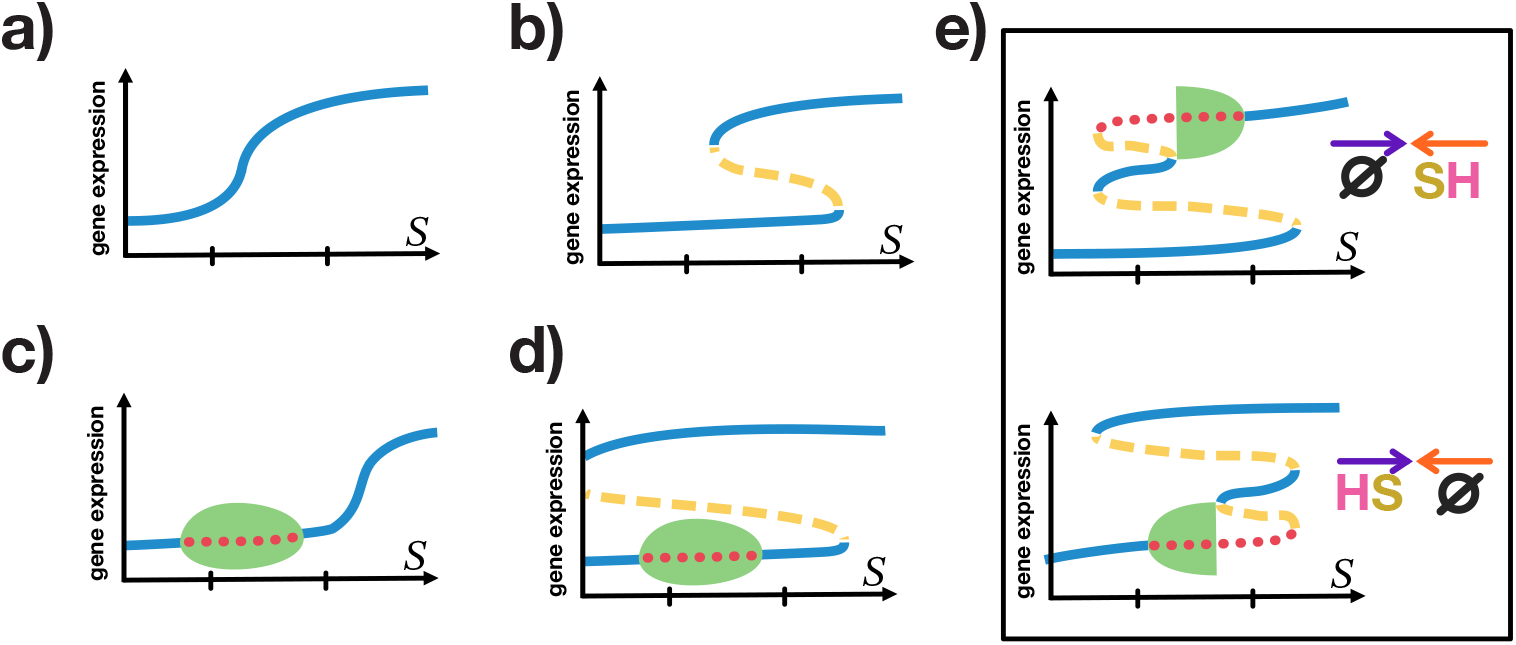
Bifurcation diagrams not part of the zoo. (a-d): Bifurcations that do not fulfil the multifunctional conditions, but that are achievable with the current model: (a) monostable behaviour, (b) bistability without oscillation, (c) oscillation without bistability, and (d) irreversible scenarios. (e) The two bifurcations that fulfil the zoo conditions, but that were not found in the search.

### 4. Bayes factor

To compare the one- and two-inducer models, we used a metric analogous to the Bayes factor, defined as the fraction of prior sampled points lying within the 90% high-density regions of the posterior KDE. We sampled the corresponding parameter spaces of 9 and 11 dimensions for the one- and two-inducer circuits, respectively, with 10^7^ points each. Using a KDE bandwidth of 0.7, 1.11×10^−5^ of points fell within the 90% region for the one-inducer circuit, compared to 8.3×10^−6^ for the two-inducer circuit. The resulting ratio (≃ 1.45) suggests that the one-inducer model has a slightly larger effective relative spread. This suggests that there is no structural disadvantage in focusing on the one-inducer circuit when designing AC-DC multifunctionality.

### 5. Machine learning classification

The 4 × 10^4^ posterior parameter sets generated from ABC-SMC were classified into three types of dynamics depending on whether they exit from oscillations via a Hopf bifurcation, a saddle-loop bifurcation or SNIC bifurcation. We trained a decision tree classifier on 1459 labelled datasets (balancing the parameter sets of each category due to overrepresentation of SNIC bifurcations cf. Fig. A3). The classifier was tested on 1460 separate datasets to assess the algorithm’s performance. The algorithms were implemented using Python’s *scikit-learn* library [56].

**FIG. A3.**
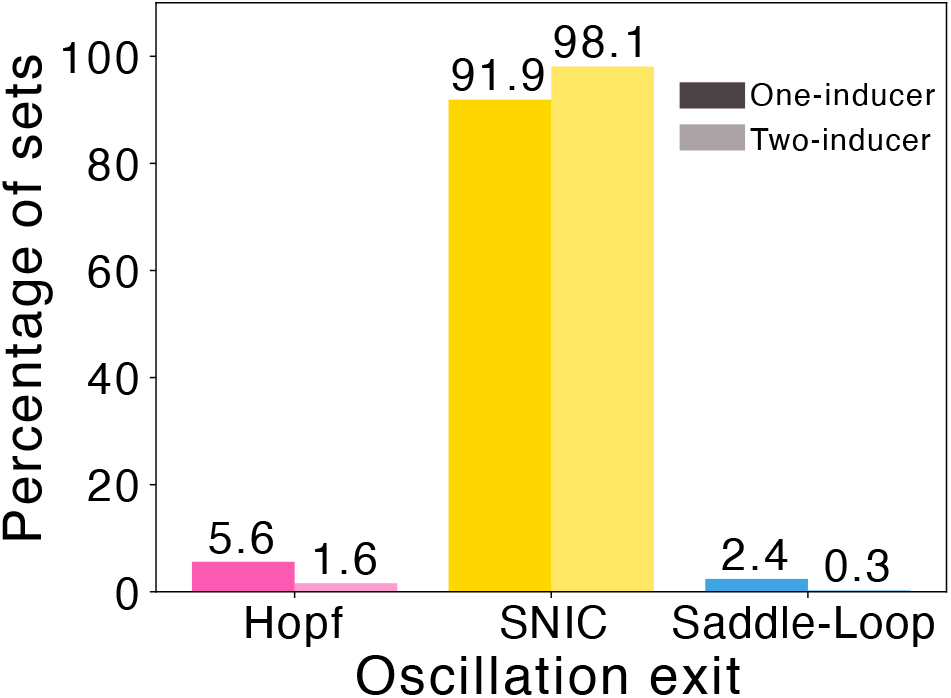
SNIC is the most prevalent exit of oscillations. Classification of exit of oscillation behaviour in the posterior sets found for one-inducer and two-inducer for the multifunctional search.

### 6. Measuring critical slowing down

To quantify critical slowing down near critical bifurcation points (e.g. at which a SNIC bifurcation or saddle-loop bifurcation) we determine the change in oscillation period near the bifurcations. For each parameter set we found the critical signal *S*_*crit*_ at which the limit cycles collide with a bifurcation point. To determine whether trajectories starting near the unstable branch have converged onto a stable branch at long times, we define an oscillation metric ℳ_*oscil*_(*S*) as the variance of gene expression *X*(*S, t*) over a specified time interval *t* ∈ [*τ*_*min*_, *τ*_*max*_]. If |ℳ_*oscil*_(*S*)| ≤ 10^−10^ we say that gene expression has converged to a stable state and no longer undergoing dynamical changes. We define *S*_*crit*_ as the minimum signal at which dynamics have converged, i.e.

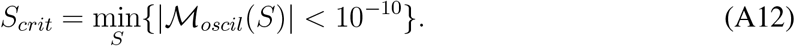

To estimate *S*_*crit*_ we evaluate ℳ_*oscil*_ over [*τ*_*min*_, *τ*_*max*_] = [198, 200]. We initially perform a coarse discretisation of the signal range *S*_*range*_ = [10, 10^3^] over 1500 points, before performing a fine-grained search over *S*_*range*_ = [*S*_*crit*_ − 20, *S*_*crit*_ + 20] with 300 points. Finally, for the signal values satisfying Δ*S* = *S*_*crit*_ − *S >* 0 we complete a fast-Fourier transform analysis with Python’s *NumPy* fft library to obtain oscillation periods *T*_*period*_. Using Python’s *SciPy* optimise we perform a linear fit to log_10_ *S* and log_10_ *T*_*period*_ over 5 points closest to *S*_*crit*_, such that

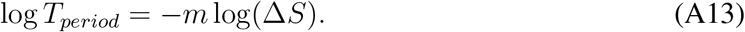

Here, *m* is the slope that quantifies the critical slowing down. Large *m* indicate significant slowing down near a bifurcation over a small signal range.

**TABLE 1.** Range of the log-uniform prior distributions used in the inference of the AC-DC model. Parameters that are only relevant in the two inducer model are indicated with ^**^. Note that log *δ*_*Y*_ = log *δ*_*Z*_ = 0 and *n* = 3.

| **Parameter** | **Lower limit** | **Upper limit** |
| --- | --- | --- |
| $\log K_{XP}$ | -5.5 | 5 |
| $\log K_{XS}$ | -5.5 | 5 |
| $\log K_{XZ}$ | 0 | 9.5 |
| $\log K_{YP}$ | -5.5 | 6 |
| $\log K_{YX}$ | -1.5 | 9.5 |
| $\log K_{ZP}$ | -5.5 | 5.5 |
| $\log K_{ZY}$ | -1.5 | 9.5 |
| $\log K_{ZX}$ | -1.5 | 9.5 |
| $\log c_{XPS}$ | 0.5 | 11.5 |
| $\log K_{YS}^{**}$ | -5.5 | 5.5 |
| $\log c_{YPS}^{**}$ | 0.5 | 11.5 |

**TABLE 2.** Parameters used in oscillator systems shown in figures 1, 3 and 4.

| Parameter | Figure 1 | Figure 3 | Figure 4 |
| --- | --- | --- | --- |
| $\log K_{XP}$ | -4.477409364333552 | -4.896113214164697 | -4.896113214164697 |
| $\log K_{XS}$ | -4.701956656436723 | -4.89651872528574 | -4.89651872528574 |
| $\log K_{XZ}$ | 1 | 1 | 1.75 |
| $\log K_{YP}$ | 2.3941003059589474 | 4.626484134323584 | 4.626484134323584 |
| $\log K_{YX}$ | 4.3131549866306775 | 5.7221227036539455 | 5.7221227036539455 |
| $\log K_{ZP}$ | 2 | [2,0.7,0.5] | 2 |
| $\log K_{ZY}$ | 4.146262028243407 | 5.220185667727906 | 5.220185667727906 |
| $\log K_{ZX}$ | 2.059934003306114 | 5.220185667727906 | 2.120148700492043 |
| $\log c_{XPS}$ | 6.128830469539324 | 6.455640251965155 | 6.455640251965155 |
| $\log \delta_Y$ | 0 | 0 | 0 |
| $\log \delta_Z$ | 0 | 0 | 0 |

**FIG. A4.**
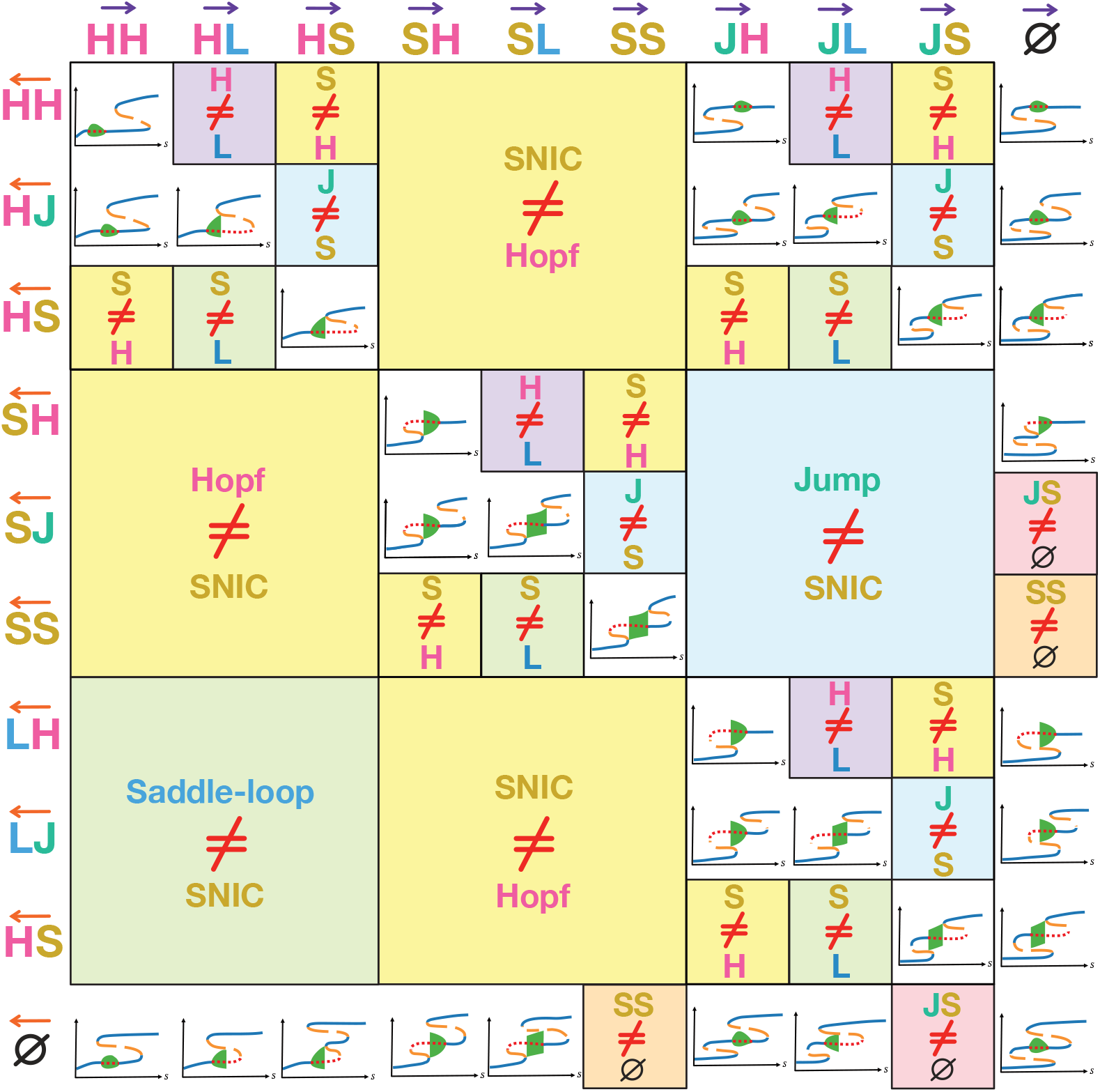
Not every barcode is geometrically achievable. Grid showing all 100 possible barcode combinations: 31 valid barcodes are illustrated with their corresponding bifurcation diagrams, while the remaining 69 infeasible ones are marked with coloured boxes indicating the reason for incompatibility.

**FIG. A5.**
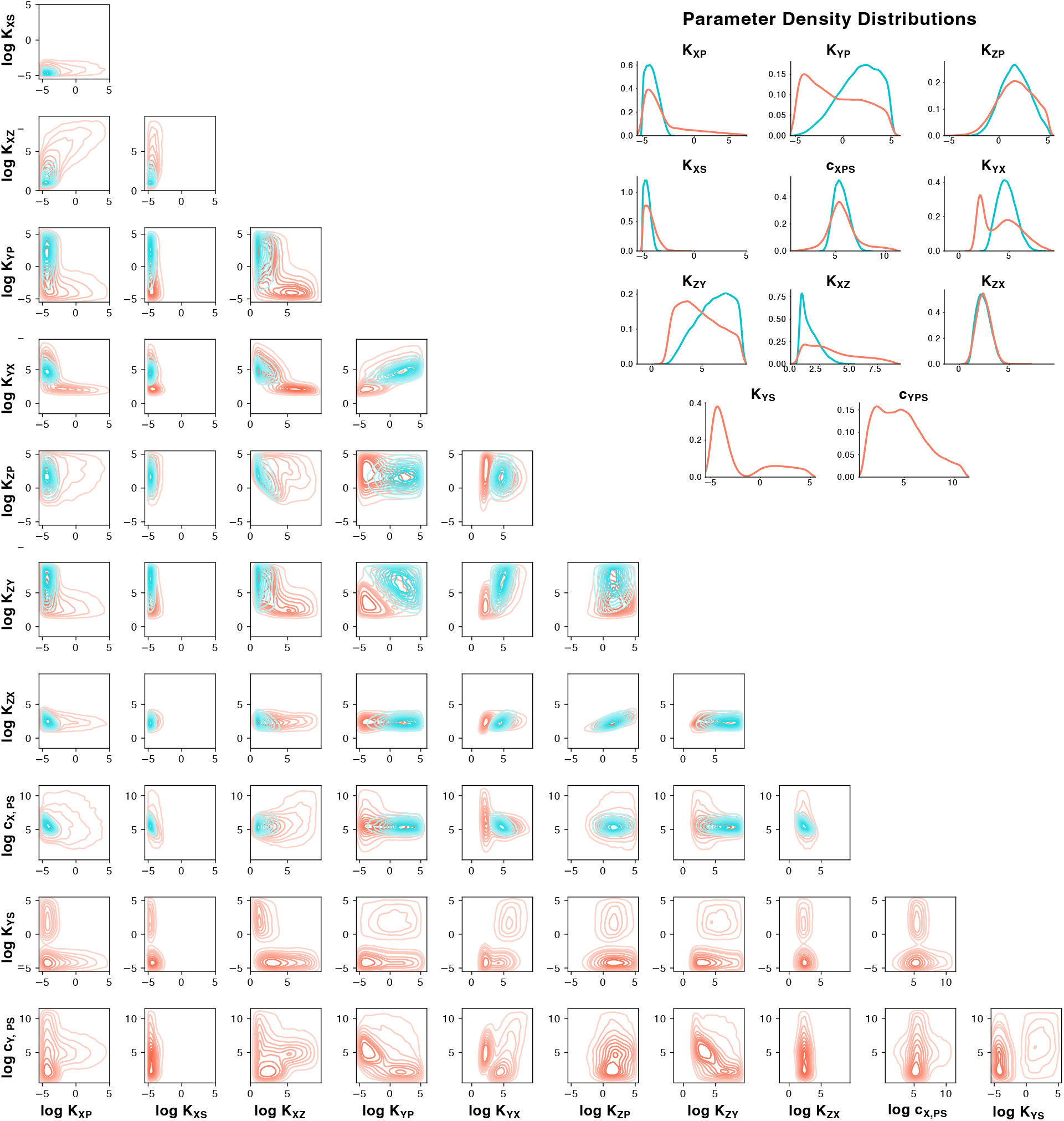
1i (blue) parameter distributions overlap with 2i (pink) parameter distributions. Bottom left: posterior probability distributions for each pair of parameters. Top right: marginal posterior distribution for the binding affinities *K*_*ji*_ and cooperativity parameters *c*_*i*_.

**FIG. A6.**
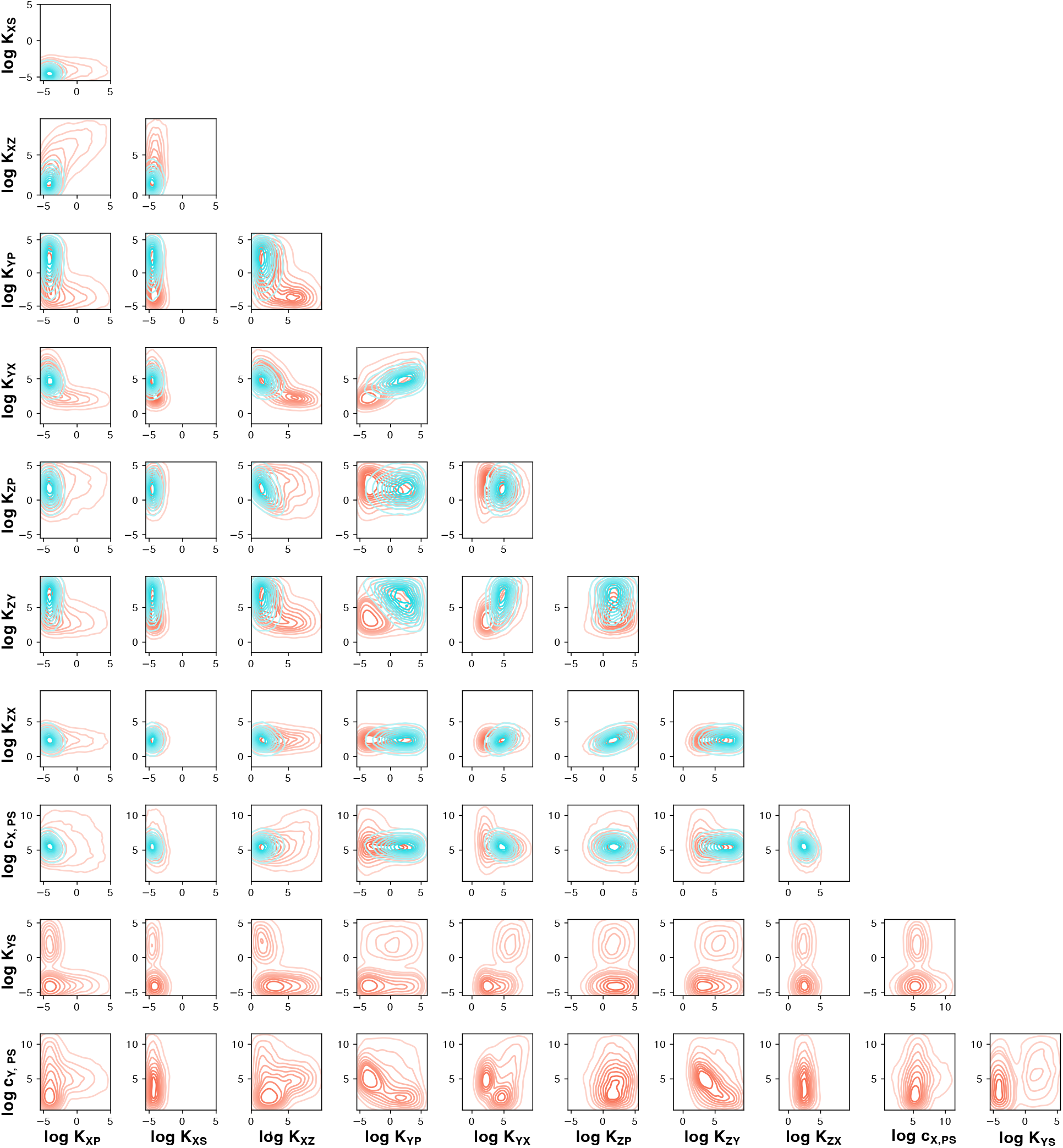
Parameter distributions resulting from resampled distributions of KDEs with bandwidth 0.7 for 1i (blue) and 2i (pink). Posterior probability distributions for each pair of parameters sampled from a 0.7 bandwidth KDE.

**FIG. A7.**
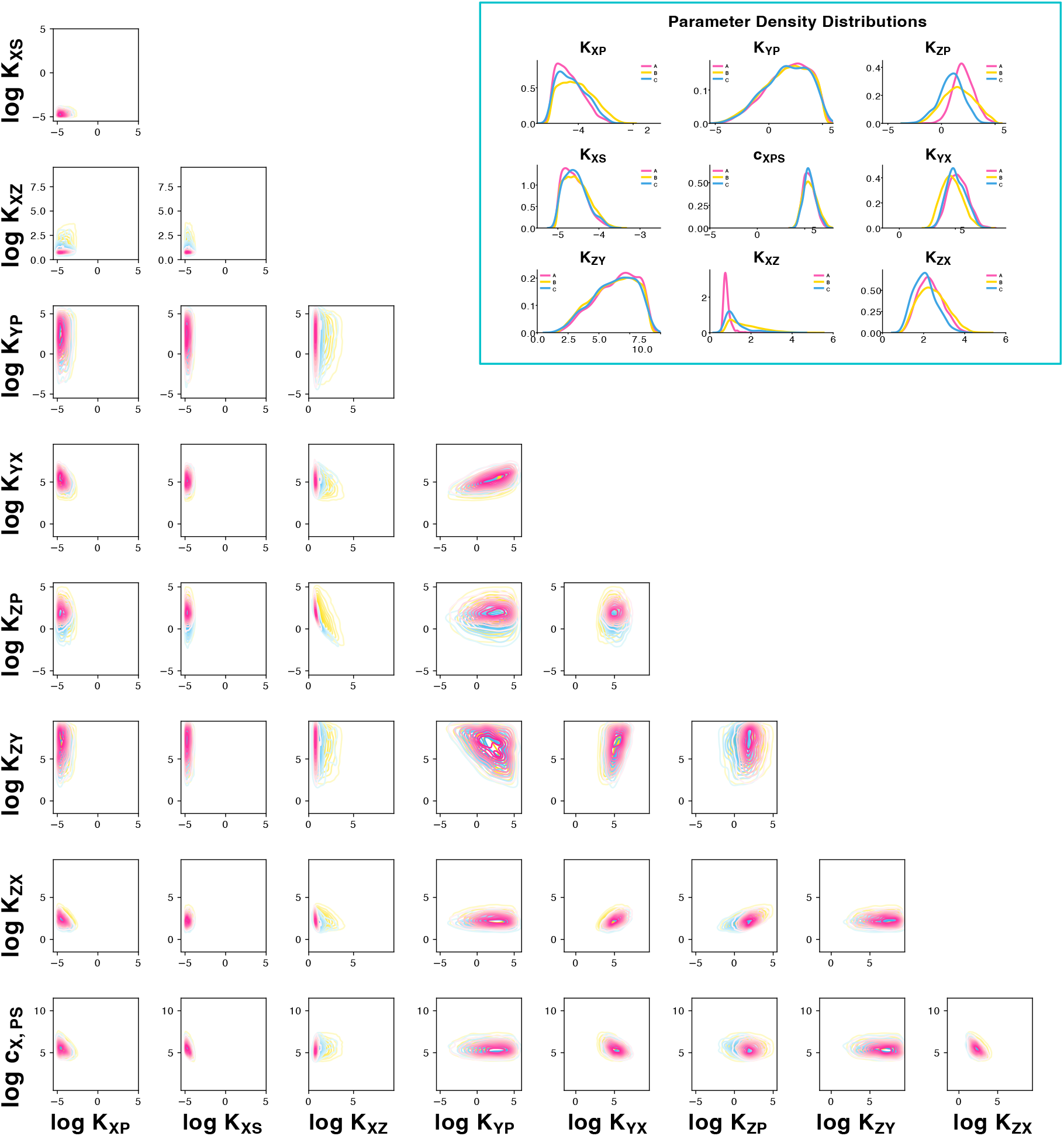
Posterior distributions for 1i categorised based on resulting bifurcation diagrams. Yellow, blue, and pink lines correspond to SNIC, Saddle-Loop and Hopf exit bifurcations, respectively. Bottom left: pairwise posterior joint distributions for each pair of parameters. Top right: marginal posterior distribution for the binding affinities *K*_*ji*_ and cooperativity parameters *c*_*i*_.

